# All-optical electrophysiology in hiPSC-derived neurons with synthetic voltage sensors

**DOI:** 10.1101/2021.01.18.427081

**Authors:** Francesca Puppo, Sanaz Sadegh, Cleber A. Trujillo, Martin Thunemann, Evan Campbell, Matthieu Vandenberghe, Xiwei Shan, Ibrahim A Akkouh, Evan W. Miller, Brenda L. Bloodgood, Gabriel A. Silva, Anders M. Dale, Gaute T. Einevoll, Srdjan Djurovic, Ole A. Andreassen, Alysson R. Muotri, Anna Devor

## Abstract

Voltage imaging and “all-optical electrophysiology” in human induced pluripotent stem cell (hiPSC)-derived neurons have opened unprecedented opportunities for high-throughput phenotyping of activity in neurons possessing unique genetic backgrounds of individual patients. While prior all-optical electrophysiology studies relied on genetically encoded voltage indicators, viral transduction of human neurons with large or multiple expression vectors can impact cell function and often lead to massive cell death. Here, we demonstrate an alternative protocol using a synthetic voltage sensor and genetically encoded optogenetic actuator that generate robust and reproducible results. We demonstrate the functionality of this method by measuring spontaneous and evoked activity in three independent hiPSC-derived neuronal cell lines with distinct genetic backgrounds.

## Introduction

Traditionally, neuronal electrical properties have been evaluated using intracellular electrophysiological recordings which are highly accurate but also terminal, invasive, low-throughput and labor intensive. Recent advances in optical microscopy and optogenetics offer the new experimental paradigm of “all-optical electrophysiology” (Hochbaum et al 2014, Kiskinis et al 2018), where genetically encoded voltage indicators (Yang & St-Pierre 2016) and optogenetic (OG) actuators (Deisseroth 2015) are combined for all-optical stimulation and readout of neuronal activity (Emiliani et al 2015). Applied biological model systems derived from human induced pluripotent stem cells (hiPSCs) (Takahashi et al 2007, Yu et al 2007), all-optical electrophysiology opens unprecedented opportunities for non-terminal and non-invasive phenotyping of neurons and neuronal networks with unique genetic backgrounds of individual patients (Brennand et al 2011, Marchetto et al 2010, Mertens et al 2015).

In prior studies, the voltage sensor and OG actuator were co-expressed using viral transduction (Bando et al 2019, Hochbaum et al 2014, Kiskinis et al 2018). However, growing hiPSC-derived neuronal networks is a challenging procedure (Calabrese et al 2019). One specific problem is that introduction of large or multiple expression vectors can often lead to high rate of cell death (Detrait et al 2002, Ding & Kilpatrick 2013, Johnston et al 2020). This, in turn, can bias the composition of the remaining neuronal networks towards cell types that are for some reason resistant against the transfection procedure. Moreover, this bias may differ across the patient and control cell lines confounding experimental results. Here, we sought to establish an alternative protocol that uses synthetic voltage sensors (Miller 2016, Peterka et al 2011, Walker et al 2020a, Walker et al 2020b) that can be easily delivered to cell cultures mitigating the problem of cell death due to introduction of large or multiple expression vectors. We demonstrated the functionality of this method by measuring spontaneous and OG-induced activity in three independent hiPSC-derived neuronal cell lines with distinct genetic backgrounds.

## Results

All-optical electrophysiology requires that the voltage sensor and OG actuator are spectrally orthogonal in order to minimize the crosstalk, i.e., avoiding excitation of the voltage sensor by light that controls the OG actuator and vice versa (Emiliani et al 2015, Hochbaum et al 2014, Kiskinis et al 2018). We chose BeRST-1 (Berkeley Red Sensor of Transmembrane potential) as the most red shifted among the available synthetic voltage indicators that can be safely delivered to all cells and offers fast response kinetics and high sensitivity to single spikes (Huang et al 2015). We combined it with photoactivation of the OG actuator CheRiff that has been previously used in all-optical electrophysiology experiments (Hochbaum et al 2014). CheRiff is a genetically encoded actuator that requires viral transduction (see **Methods**); however, the impact of transduction with lentiviruses carrying a CheRiff-EGFP expression cassette alone on cell viability was not significant in our hands. We also co-loaded cells with a synthetic calcium (Ca^2+^) indicator Oregon Green BAPTA-1 AM (OGB1) to perform Ca^2+^ imaging prior to voltage imaging to identify active fields of view (FOVs).

### Optimization of imaging protocol

The imaging setup consisted of a body of an inverted epifluorescence microscope with a custom illumination light path (**Fig. 1A**). The illumination was provided by a pair of diode lasers at 635-nm and 473-nm for simultaneous BeRST-1 imaging and CheRiff actuation or, alternatively, sequential imaging of BeRST-1 and OGB1 (**Fig. 1B**) (see **Methods**). The laser beams were collimated and combined by a dichroic mirror. To reduce background fluorescence due to out-of-focus excitation we implemented a custom solution for oblique angle illumination (Werley et al 2017) significantly improving the signal-to-background ratio (SBR) (see **Methods**, **Fig. 1C**).

**Figure 1.**
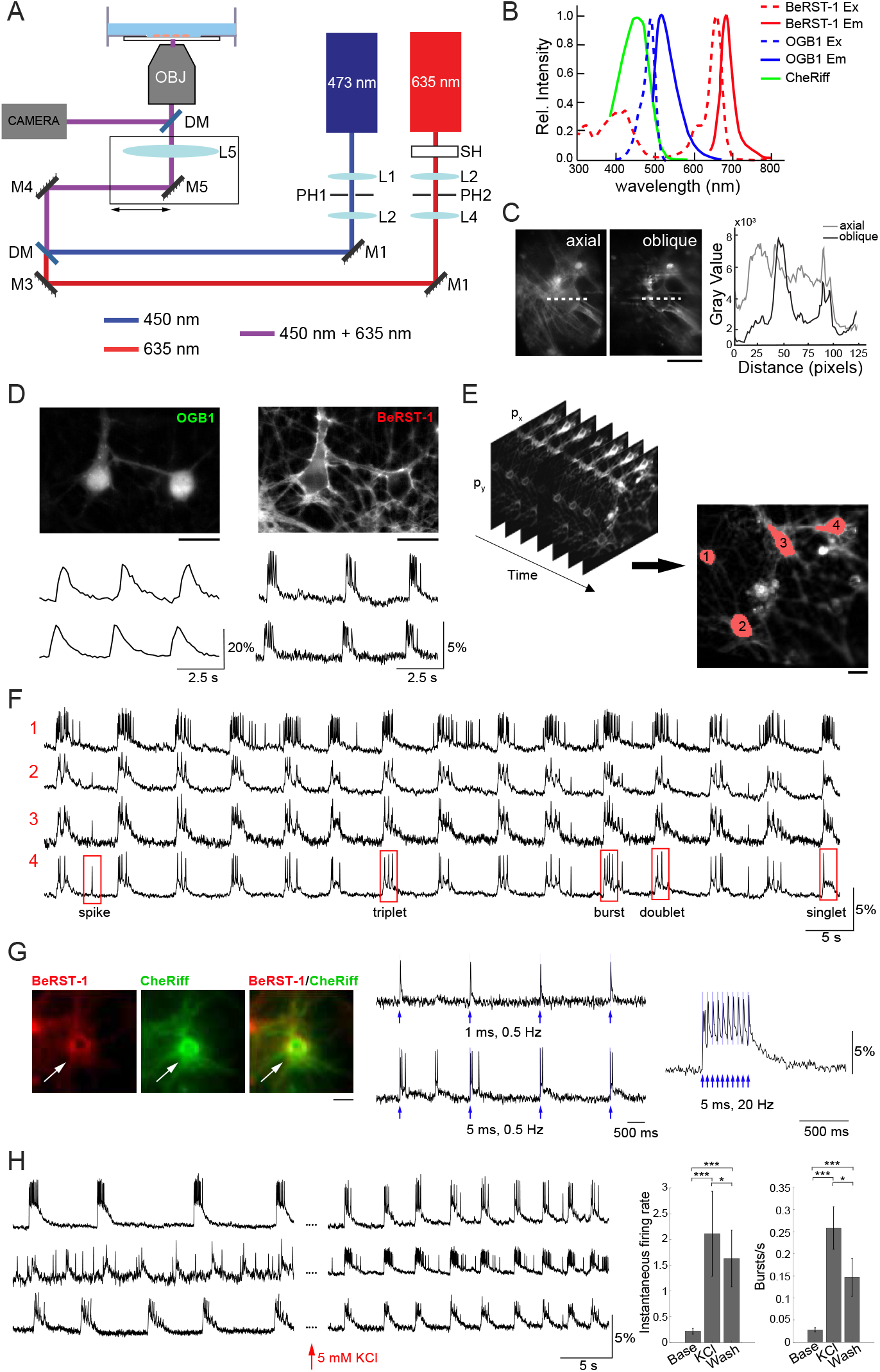
Optimization of the imaging protocol in primary neurons. **A.** Imaging Setup. The 635-nm and 473-nm laser beams were collimated and spatially filtered through a combination of lenses (L1-L4) and pinholes (PH1 and PH2). A shutter SH was used to block the 635-nm beam when not acquiring data; the 473-nm beam was modulated using an analog signal. The beams were combined by a dichroic mirror DM1 and focused onto the rear focal plane of the objective via a lens LS. The lens and mirror M5 were translated together in the plane orthogonal to the optical axis to offset the illumination enter the objective producing oblique illumination. Fluorescence was directed to a camera for detection of BeRST-1 or OGB1 signals through a dichroic mirror an emission filter. **B.** Overlaid excitation (dashed line) and emission (solid line) spectra of BeRST-1 (red) (Huang et al 2015) and OGB1 (blue, reproduced from Fisher Scientific) and action spectrum of CheRiff (green) (Hochbaum et al 2014). **C.** Comparison of BeRST-1 fluorescence profile with axial and oblique illumination (gray and black lines, respectively). Scale bar, 10 μm. **D.** Primary neurons co-labeled with OGB1 (left) and BeRST-1 (right). Scale bar, 10 μm. Time-courses of spontaneous Ca^2+^ and voltage activity extracted from these two neurons are shown below the respective OGB1 and BeRST-1 images. **E.** Segmentation of single-neuron ROIs. Scale bar, 10 μm. **F.** Extraction of single-neuron voltage time-courses. The four traces in (F) correspond to four different neurons segmented in (E). **G.** All-optical electrophysiology with BeRST-1 in CheRiff-expressing neurons. Left: Primary neurons stained with BeRST-1 (red) and expressing CheRiff-EGFP (green). Right: Voltage response to OG stimulation of varying frequency and duration. **H.** High K^+^ increases spiking activity in primary neurons. Left: Voltage time-courses of three representative neurons in normal imaging buffer and after perfusion with 5 mM KCl. Right: Instantaneous firing rate and bursting rate at baseline (Base, n = 9 neurons), after 5 min of perfusion with 5 mM KCl (KCl, n= 9 neurons) and after washing off high K^+^ (Wash; n = 25). Error bars indicate mean ± SD; unpaired Student’s t-test (*p < 0.05; **p < 0.01; ***p< 0.001).

To troubleshoot our imaging protocol, we used primary cultures of rat dissociated hippocampal neurons at 14-23 days in vitro (DIV) that had robust and synchronous spiking activity (see **Methods** and **Figure 1 - figure supplement 1A-F**). Co-labeling with OGB1 was critical for quick and efficient evaluation of the level of spiking activity and for choosing FOVs for subsequent voltage imaging. Further, cytosolic OGB1 staining facilitated visual inspection of the cell culture including cell morphology, density, and connectivity (**Fig. 1D**). We imaged spontaneous voltage activity continuously for ~3 min using FOVs of ~ 300 × 150 μm acquired at ~500 Hz, (2.2 ms exposure, 25 mW/cm^2^ laser power). Consecutive acquisition periods were separated by at least 2 min where the 635-nm illumination was blocked by a mechanical shutter. Under this regime, we observed virtually no photobleaching (see **Figure 1 - figure supplement 1G**). The size of FOV was limited by the camera rate. With a 20x objective, our FOVs included up to 12 neurons. Image analysis and segmentation were adapted from published methods (Hochbaum et al 2014, Mukamel et al 2009). The algorithm was used to extract several key parameters describing firing properties of individual neurons including the shape of the action potential (AP), number of spikes, plateau depolarizations, and different types of bursting behavior (**Fig. 1E-F** and **Methods**). Primary neurons at 17 DIV had AP duration of 10.1±7.14 ms (mean ± SD; n=27 neurons, 635 APs) at 30 °C in the imaging chamber (see **Methods** and **Figure 1 - figure supplement 2**).

Next, we transducted primary neurons with lentivirus carrying CheRiff-EGFP expression cassette under the CamKIIα promoter for all-optical electrophysiology (**Fig. 1G**). BeRST-1 has a second, smaller excitation peak around 420 nm. In our hands, OG stimulation at 450 nm (that is commonly used for excitation of blue OG actuators) resulted in excitation of BeRST-1 seen as an increase in the voltage signal following the shape of the 450-nm laser pulse (not shown). Switching to a 473-nm laser eliminated this cross talk. For troubleshooting the protocol, we used 5 mM potassium chloride or 50 μM picrotoxin (gamma aminobutyric acid antagonist) as a means for inducing high-level spiking activity (**Fig. 1H** and **Figure 1 - figure supplement 3**). This procedure was used to evaluate the ability of neurons to produce spikes in viable neurons “on demand” during the troubleshooting of the protocol irrespective of the level of CheRiff expression.

### Optimization of hiPSC-derived cell culture protocol

Next, we translated our imaging protocol to hiPSC-derived neuronal cell cultures. The success rate of our experiments critically depended on obtaining healthy, active, and spatially uniform monolayer cultures of differentiated neurons (**Fig. 2A**). To this end, we modified a previously published procedure (Calabrese et al 2019) aiming to promote the growth of healthy, well-adhered, sparse monolayer cultures with limited clustering and minimal presence of dead cells (see **Methods** and **Figure 2 - figure supplement 1**). Our protocol started with the initial 4-5 weeks of neural progenitor cells (NPC) differentiation followed by replating and further 3-4 weeks of differentiation before transduction with the lentivirus carrying CheRiff-EGFP (see **Methods** and **Figure 2 - figure supplement 1**). Compared to the protocol of Calabrese et al. (Calabrese et al 2019), we delayed the replating procedure to prevent proliferation of undifferentiated cells and improved the dissociation technique (see **Methods**). All-optical electrophysiology was performed 7 to 10 days after the viral transduction when robust EGFP florescence indicated successful expression of the transgene (**Figure 2 - figure supplement 1**).

**Figure 2.**
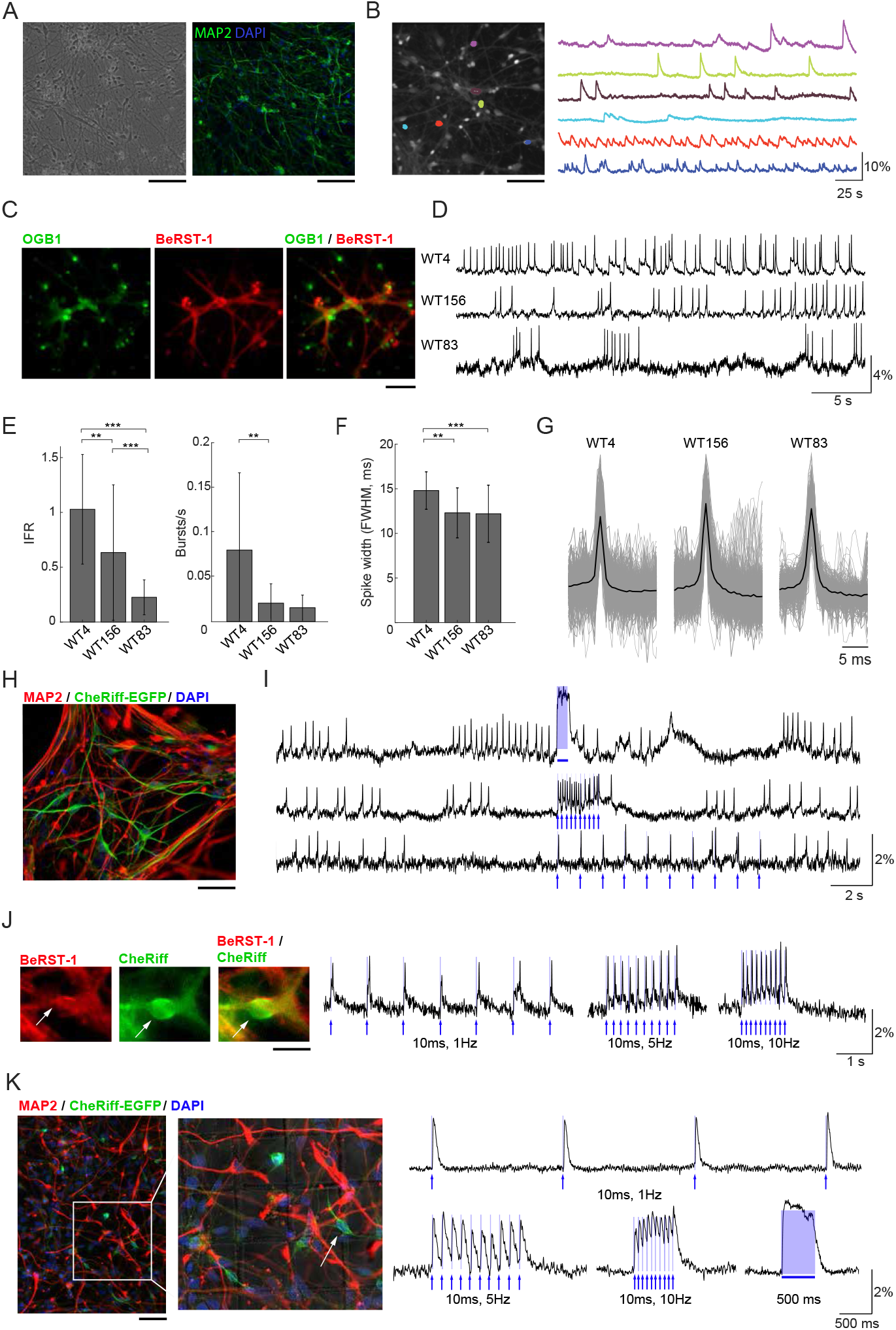
Voltage imaging and all-optical electrophysiology in human neurons. **A.** Monolayer cultures of human neurons. Left: Transmitted light image of cell culture after 8 weeks of differentiation. Right: cell culture after 10 weeks of differentiation immunostained for MAP2 and DAPI. Scale bars, 200 μm. **B.** Ca^2+^ imaging in human neurons. Left: A representative FOV with OGB1-loaded neurons. Right: Color-coded Ca^2+^ time-courses from six ROIs corresponding to individulal neurons. Scale bar, 200 μm. **C.** Human neurons stained with OGB1 (left, green), BeRST-1 (middle, red), and overlay of the two signals (right). **D.** Voltage imaging in human neurons with BeRST-1. Representative time-courses of spontaneous voltage actvity in three hiPSC-derived control cell lines showing different levels of activity and distinct firing patterns. **E.** Quantification of instantaneous firing rate and bursting rate across the three cell lines (n = 24 neurons in line WT4; n = 36 in line WT156; n = 41 in line WT83). Data are shown as mean ± SD; unpaired Student’s t-test. **F.** AP waveforms for the three control cell lines (line WT4: n = 16 neurons, 383 APs; line WT156: n = 12, 2349 APs; line WT83: n = 8, 783 APs. **G.** Distribution of the AP duration (full width at half maximal (FWHM) amplitude) for the APs shown in (E): line WT4, 14.8 ± 2.1 ms; line WT156, 12.3 ± 2.8 ms; line WT83, 12.2 ± 3.2 ms. Data are shown as mean ± SD. **H.** Immunostaining of human neurons for MAP2 (red) and EGFP (green). The nuclei were counterstained with DAPI (blue). Scale bar, 350 μm. **I.** OG stimulation in spontaneously active CheRiff-expressing control neurons (line WT4). The top trace shows activity evoked by stimulation with a continuous 500-ms OG stimulus; the two bottom traces show stimulation with 5-ms long light pulses of different frequencies (1 and 10 Hz). The timing of OG stimulation is indicated in blue. **J.** Evoked depolarization and spiking in human neurons with low spontaneous activity. Left: A representative human neuron expressing CheRiff-EGFP (green) and stained with BeRST-1 (red). Scale bar, 15 μm. Right: spiking induced by 10-ms long OG stimulation of increasing frequency (1, 5 and 10 Hz). **K.** Left: Post-hoc immunolabeling with MAP2 (red), and EGFP (green); the nuclei are counterstained with DAPI (blue). The localization grid is visible in the zoomed-in image. Scale bar, 500 μm. Right: all-optical electrophysiology from one neuron prior to fixation. The timing of OG stimulation is indicated in blue.

### Voltage imaging in hiPSC derived neurons with BeRST-1

Similar to primary neurons in **Figure 1**, we used OGB1 for quick evaluation of the level of activity in human neurons (**Fig. 2B**) taking advantage of larger FOVs achievable with Ca^2+^ imaging due to a tradeoff between FOV and imaging speed (Kulkarni & Miller 2017). Co-labeling of hiPSC-derived cultures with BeRST-1 and OGB1 (**Fig. 2C**) was also used for visual inspection for remaining undifferentiated NPCs or overgrowth of glia providing feedback for optimization of the cell culture protocol (see **Methods** and **Figure 2 - figure supplement 1**). We applied our protocol to three independent hiPSC-derived neuronal cell lines with distinct genetic backgrounds (WT4, WT83, and WT156). In agreement with previous reports (Paavilainen et al 2018, Shi et al 2012), spontaneous activity was low and heterogeneous during early development increasing after ~8 weeks of differentiation (withdrawal of basic fibroblast growth factor (bFGF) from the medium, see **Methods**). Interestingly, we observed clear differences in the firing patterns between the three cell lines (**Fig. 2D-E**). Most noticeably, the firing rate varied between 1.03±0.50-Hz in line WT4 (n=24 neurons) to 0.23±0.16-Hz in line WT83 (n=41 neurons) (**Fig. 2E**). The AP waveform did not differ significantly across the cell lines (**Fig. 2F-G**).

Next, we transducted human neurons with lentivirus carrying CheRiff-EGFP expression cassette under the CamKIIα promoter for all-optical electrophysiology (**Fig. 2H**). We observed higher heterogeneity in EGFP fluorescence in human neurons compared to primary neurons, likely reflecting variable levels of CheRiff-EGFP expression in human neurons at different stages of maturation. Photoactivation of CheRiff robustly induced depolarization and spiking irrespective of the level of spontaneous activity (**Fig. 2 I-J**).

The heterogeneity of firing properties within a cell line, as indicated by the error bars in **Figure 2E**, could be at least in part due to differences in neuronal cell types. In contrast to genetically encoded voltage sensors, synthetic probes such as BeRST-1 cannot be targeted to specific cell types. This limitation, however, can be mitigated by fixation and post-hoc immunolabeling, as long as the fixation and labeling procedure does not distort the cellular network allowing for registration with live images. Post hoc labeling and image registration have been performed on cell cultures in the past (Hanson et al 2010, Walker et al 2020b) but not in the context of voltage imaging in human neurons. As a proof of principle, we plated neurons in imaging dishes with an imprinted grid. Individual neurons were targeted for OG stimulation and voltage imaging following by *in situ* fixation and immunolabeling immediately after the imaging session (see **Methods**). With this procedure, we were able to find FOVs and specific neurons used for live imaging (**Fig. 2K**). Although the majority (~90%) of neurons produced with our culturing protocol were glutamatergic (Marchetto et al 2017, Zhang et al 2016), in the future, this protocol may help addressing cell-type-specific neuronal activity while circumventing the need for genetic encoding of voltage sensors.

## Conclusions

In summary, voltage imaging with synthetic probes such as BeRST-1 combined with genetically encoded OG actuators provides a robust paradigm for all-optical electrophysiology in hiPSC-derived neurons and neuronal networks possessing unique genetic background of individual patients. In the present study, this protocol has proven to be very effective offering high quality imaging readouts while mitigating cell death due to introduction of large or multiple expression vectors. While synthetic probes such as BeRST-1 indiscriminately label all membranes, the cell type of imaged neurons can be identified post hoc using immunostaining and image registration. The present observation of varying firing patterns across the considered control cell lines supports the use of isogenic controls whenever possible, as previously suggested (Iakoucheva et al 2019). We envision that the same protocol, including fixation after all-optical-electrophysiology, will be used in future studies for single-cell-resolved nuclei-isolated RNAseq or *in situ* RNA hybridization from imaged neurons providing a link from the functional phenotype to the underlying gene expression profile.

## Methods

### Imaging setup

We engineered our imaging setup around a body of an inverted Olympus IX71 epifluorescence microscope. Illumination was provided by a pair of CW diode lasers at 635-nm (500mW, Opto Engine LLC) and 473-nm (100mW, Cobolt 06-MLD). The laser beams were collimated, spatially filtered, and combined by a dichroic mirror (Semrock, FF01-370/36-25). After the mirror, co-aligned beams were directed towards a lens (AC508-400-A-ML, f = 400 mm) focusing the light onto the rear focal plane of a high numerical aperture objective (Olympus UPlanFL N 20x, numerical aperture (NA) = 0.5 (air) or Olympus UPlanFL N 40X/NA = 1.30 (oil)). The lens (and a mirror in front of the lens) was translated in the plane orthogonal to the optical axis displacing the focal spot off the axis in the rear focal plane of the objective. This produced oblique illumination, where the laser light propagated at a low angle close to the bottom of the culture dish after refracting at the glass-water interface (Werley et al 2017). By restricting the fluorescence to a region of the specimen near the bottom of the dish, oblique illumination reduced the background improving the signal-to-background ratio (SBR). The angle of incidence of the light onto the specimen depended of the amount of offset of the lens allowing fine adjustment for achieving the best results. Oblique illumination was in particular important in cases with overlapping cells and out-of-focus debris.

The 635-nm beam was used for BeRST-1 imaging with intensity at the sample of 25 W/cm^2^. A shutter was placed in front of the 635-nm beam to control the illumination time of the sample without the need for turning the laser off, which would lead to power instability. The 473-nm beam was used for CheRiff actuation with intensity at the sample of 5-10 mW/cm^2^. Laser power of the 473-nm beam was controlled by an analog square signal provided by a computer-controlled National Instruments Data Acquisition (DAQ) board. The same 473-nm beam was used for OGB1 imaging with intensity at the sample of 8 mW/cm^2^ in experiments that did not involve OG stimulation.

BeRST-1 fluorescence was filtered at 736±64 nm and collected using a scientific sCMOS camera (Zyla 4.2 Plus, Andor) operated at 300-500 Hz, 2.2 ms exposure, 1x gain, FOV of 200 x 150 μm. OGB1 fluorescence was filtered at 535±25 nm and collected with the same camera operated at 50 Hz. The emission filters were swapped in between voltage and Ca^2+^ imaging. Data were collected in epochs of ~3 min for imaging of spontaneous neuronal activity and 30 s for all-optical electrophysiology. Consecutive data acquisition epochs were separated by at least 2 min where the 635-nm illumination was shuttered off.

### Primary neuronal cultures

All animal procedures were performed in accordance with the University of California San Diego Institutional Animal Care and Use Committee and complied with all relevant ethical regulations for animal research. Dissociated hippocampal neuronal cultures were derived from wild-type Sprague Dawley rat pups of both sexes at postnatal days P0-P1. The cell cultures were prepared as previously described (Goslin et al 1991), and plated at a density of 130 cells per mm^2^ on laminin-coated 35-mm MatTek dishes (MatTek CatP35G-0-14-C). Neurons were grown in Neurobasal-A media (ThermoFisher Scientific Cat10888022) supplemented with Glutamax (ThermoFisher Scientific Cat35050061), Pen/Strep (ThermoFisher Scientific Cat10378016), and B27 supplement (ThermoFisher Scientific Cat17504044).

### hiPSC-derived neuronal cultures

All iPSC were provided by the Muotri lab. Cortical NPCs were differentiated from hiPSC as described previously (Griesi-Oliveira et al 2015). To differentiate NPCs to cortical neurons, NPCs were dissociated with Accutase (StemCell Technologies), plated on a 10-cm tissue culture dishes coated with 10 μg/ml poly-L-ornithine (Sigma, P3655) and 2.5 μg/ml mouse laminin (1 mg, Invitrogen 23017-015), and then cultured for 2 weeks in NPC medium (DMEM/F12 supplemented with 1% penicillin-streptomycin, N2 NeuroPlex (Gemini Bioproducts), NeuroCult SM1 (Stem-Cell technologies) and 20 ng/mL basic fibroblast growth factor (bFGF; Life Technologies). When NPCs reached 90% confluency, NPCs were differentiated into cortical neurons by bFGF withdrawal. Differentiating cells were fed with NPC medium every 3 days. hiPSC-derived cultures were differentiated for 4-5 weeks to allow growth of healthy neurons, extension of long processes, and formation of densely interconnected networks. Neurons were then dissociated and replated on 35-mm imaging plates with glass bottoms, as described below, and kept differentiating for a further 3-4 weeks with excellent cell survival, cell recovery and connectivity re-growth.

### Cell dissociation and replating

To dissociate and replate mature neurons that had already established long neurites and a complex connectivity network, cells were dissociated with Accutase (StemCell Technologies). After 40 minutes of incubation, the dense network of cells that had lifted from the plate was mechanically dissociated by slowly pipetting up and down using a 5-ml wide-tipped Pasteur pipette. The culture was then incubated for an additional 10 minutes. Next, regular phosphate-buffered saline (PBS) was added to the plate to block the enzymatic reaction and further gentle mechanical trituration was carried out, first with a 1000-μl pipette and then with a 200-μl pipette to gradually loose and dissociate the thick networks of neurites and cell clusters. The dissociated culture was strained through a 70-μm cell strainer to remove clumps, and then centrifuged for 4 min at 900 rpm. The resulting cell pellet was re-suspended in Media2 (Neurobasal media (Life Technologies) supplemented with GlutaMAX (Life Technologies), 1% Gem21 NeuroPlex, 1% MEM nonessential amino acids (NEAA, Life technologies), and 1% Penicillin streptomycin) supplemented with 1% fetal bovine serum (FBS, Gemini Bio-products), 1 μg/ml of laminin and 0.1% ROCK (Rho kinase) inhibitor (Y-27632, Fisher Scientific). Cells were replated on a 14-mm cover slip of a 35-mm imaging plates coated with 100 μg/ml poly-L-ornithine and 5 μg/ml laminin to reach a final density of 0.3 million cells, with a typical cell viability of 80-90%. After 6 hours, additional M2 medium was added to allow dead cells to lift and be aspirated away. The neurons were then fed with fresh M2 medium supplemented with 5 μM cytosine arabinoside (AraC) (Sigma-Aldrich) to prevent proliferation of glial cells underneath the neurons. After 2 days of AraC exposure, the cells were fed with fresh M2 medium; feeding was repeated once per week with M2 medium (half media exchange).

### Gene delivery

CheRiff was delivered via lentiviral transduction after transferring to the imaging dish. Lentivirus was produced by VectorBuilder (Cyagen) from CamKIIα-CheRiff plasmid (Hochbaum et al 2014). DRH313: FCK-CheRiff-eGFP was a gift from Adam Cohen (Addgene plasmid #51693; http://n2t.net/addgene:51693; RRID:Addgene_51693). Primary neurons were transduced at 6 DIV and used for experiments at 14-23 DIV. hiPSC-derived neurons were transduced 7-10 days prior to imaging. On the transfection day, 4-12 μL of the CheRiff lentivirus were combined with 200 μL of conditioned medium from the culture to allow the cells benefit from their secreted growth factors (Yang et al 2017). Following an overnight incubation, an additional 2 ml of M2 medium were added. After one more day of incubation, the plates were washed and replenished with fresh M2 medium. hiPSC-derived neurons were fed with antibiotic-free M2 medium for one week prior to transduction, which resulted in higher expression levels and lower cell death.

### Loading of optical probes

BeRST-1 was produced as previously described (Huang et al 2015) and stored as 5 mM in DMSO. OGB1-AM (O-6807, Invitrogen, 50 μg) was first dissolved in 4 μl of 20% pluronic in DMSO (F-127, Invitrogen); 80 μl of imaging buffer (MgCl_2_ 1 mM, NaCl 130 mM, KCl 3 mM, CaCl_2_ 1 mM, Glucose 10 mM, HEPES 10 mM; pH 7.4) were added to yield the final concentration of 0.5 mM OGB1-AM. Both primary and hiPSC-derived cultures were incubated in the imaging buffer with the final concentration of 5 μM BeRST-1 and 5 μM OGB1-AM for ~20 min immediately prior to the imaging session. We did not use OGB1 in experiments involving OG stimulation. During this time, cells were kept within an incubator at 37 °C. Next, cells were washed and replenished with fresh imaging buffer. Data were acquired at 30 °C. Temperature was maintained by continuous perfusion with a heated imaging buffer (see below). Careful temperature maintenance was important, because it affected the level of activity, firing pattern, and the AP waveform (**Figure 1 - figure supplement 2**).

### Perfusion system

Cultures were perfused with imaging buffer for the entire duration of the experiment. We used a diamond-shaped bath (Warner Instruments LLC) incorporated into the imaging dish to provide a laminar flow across the culture. The speed of perfusion was controlled with a peristaltic pump (Peri-Star, WPI) at 2 ml/min. An in-line heater (SC-20, Warner Instruments) was positioned in close proximity of the imaging chamber to maintain the temperature of the imaging buffer within the dish at 30 °C. The heater was calibrated with a thermo-probe inserted into the dish. The imaging buffer containing high K^+^ (5 mM) or picrotoxin (50 μM) were washed in using computer-controlled microfluidic valves (VC-6, Warner Instruments).

### Image processing and signal analysis

The protocol for image analysis was adapted from Mukamel et al. 2009 (Mukamel et al 2009) and Hochbaum et al., 2014 (Hochbaum et al 2014). In brief, images were filtered in time and space to improve the signal-to-noise ratio (SNR). Then, a combination of principal component analysis (PCA) and independent component analysis (ICA) was used to identify regions of interest (ROIs) corresponding to individual neurons and extract corresponding single-neuron voltage time-courses. The time-courses were computed as percent change relative to the baseline (ΔF/F_0_). Single-neuron ΔF/F_0_ time-courses were computed as described in Hochbaum et al., 2014 (Hochbaum et al 2014).

For spike detection, we adapted an algorithm based on a nonlinear energy operator (NEO) that was previously used for extracellular electrophysiological recording to emphasize spike-like signals (Mukhopadhyay & Ray 1998). The NEO signal was calculated as s(t)^2^ - s(t-1) * s(t+1), where s(t) is single-neuron voltage time-course ΔF/F_0_. Peaks were identified in the NEO signal using a MATLAB peak-seek routine. All peaks below a 3 standard deviations of the NEO signal were discarded. To avoid double counting when the same AP crossed the threshold twice due to high frequency noise in the recording, we set a hard limit of <20 ms on the time between two spikes (inter-spike-interval, ISI).

To obtain AP waveforms, we extracted segments of voltage traces in a window of 300 ms centered on a detected AP peak. The waveforms were then aligned on the peak, and the waveform parameters were extracted as previously described (Bean 2007, Trainito et al 2019). The spike duration was quantified by calculating the full width at half-maximal amplitude (FWHM).

We used a custom MATLAB routine to distinguish individual APs from those riding on top of plateau depolarizations including one (singlet), two (doublet), and three (triplet) APs. A burst was defined as any sequence of four or more APs with mean inter-spike-interval (ISI) shorter than 250 ms (Chen et al). The algorithm computed instantaneous firing frequency, bursting rate, ISI, inter-burst-interval (IBI), and burst duration.

### Post hoc immunolabeling

Neurons were replated on 50-μm grids patterned on the bottom of 35-mm MatTek dishes (MatTek CatP35G-0-14-C) previously coated with 100 μg/ml poly-L-ornithine and 5 μg/ml laminin. Immediately after live imaging, cells were fixed *in situ* with 4% paraformaldehyde (PFA, Core Bio Services) for 30 minutes at room temperature. The cells were then washed 3 times in phosphate buffered saline (PBS), permeabilized with 0.2% Triton X-100 in PBS for 15 minutes and blocked with 2% bovine serum albumin (BSA, Gemini Bio-products) in PBS for 2 hours at room temperature. Primary antibodies (chicken anti-MAP2, Abcam ab5392, 1:500; rabbit anti-GFP, Molecular Probes A-21311, 1:1000) were diluted in blocking buffer (2% BSA in PBS). Cells were incubated with the primary antibodies overnight at 4 °C. The following day cells were washed 3 times in PBS, and incubated with the secondary antibodies coupled to Alexa Fluor 488, 555, and 647 (Life Technologies, 1:1000 in blocking buffer) for 1 hour at room temperature. Cells were then washed 3 times in PBS to remove the secondary antibodies, and DAPI (1:10,000 in PBS) was added for 30 minutes at room temperature to counterstain cell nuclei. Cells were then washed again 3 times in PBS. Finally, the PBS was aspirated, and the cells were submerged in a drop of ProLong Gold anti-fade mountant (Life Technologies) and covered with a glass coverslip for subsequent analysis.

### Statistics

All results are expressed as the mean ± standard deviation from mean. For evaluation of statistical significance, datasets were compared using unpaired 2-tailed Student’s t test. p values < 0.05 were considered significant.

## Supplementary Information

**Figure 1 - figure supplement 1.**
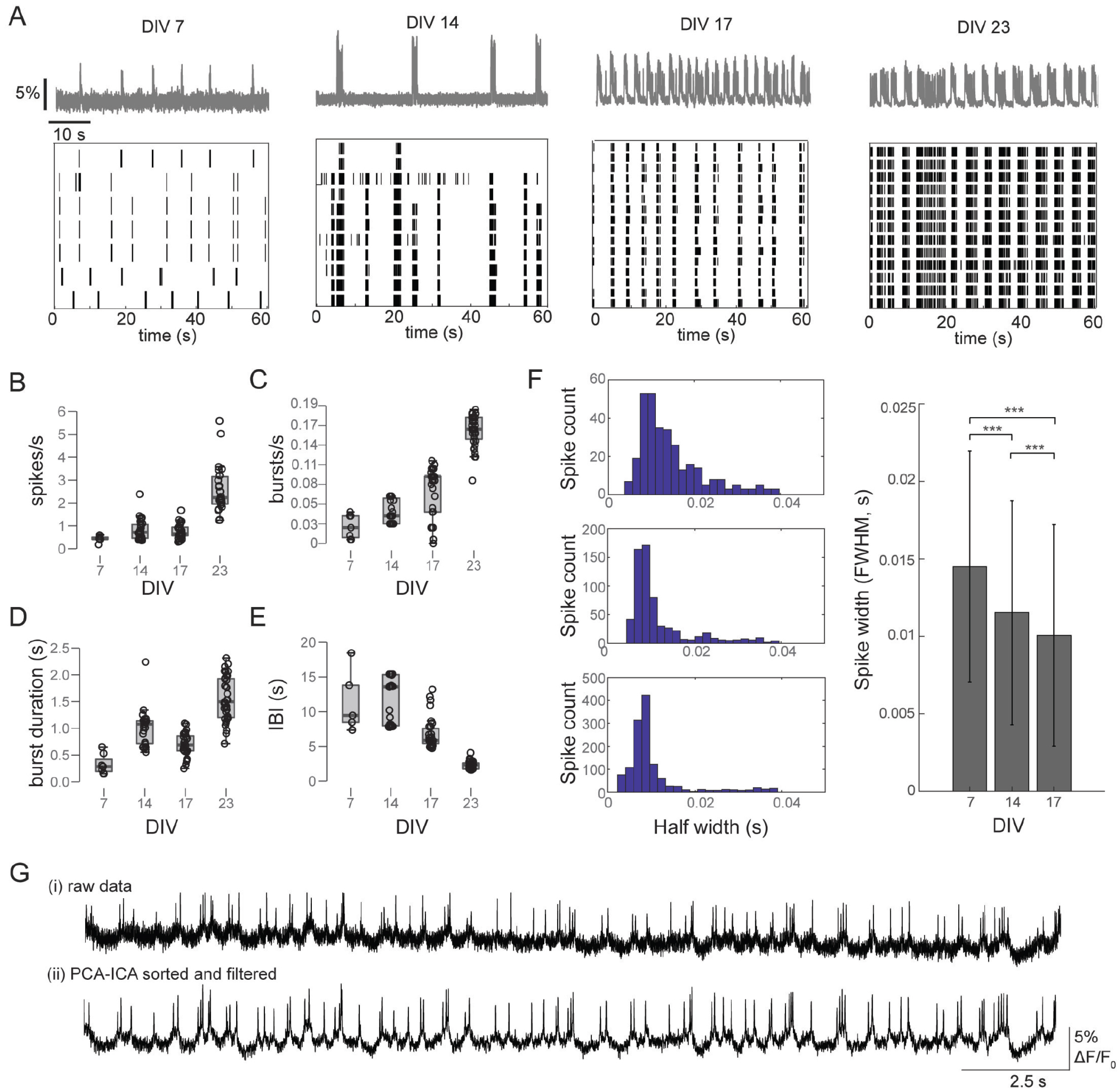
Voltage imaging across developmental stages in primary neurons. **A.** In primary neurons, the level of activity and degree of network synchronization increased from DIV 7 to DIV 23. This was accompanied by the emergence of network bursts. Top: Representative voltage time-courses recorded at 7, 14, 17 and 23 DIV. Bottom: raster plots showing spike timing for a subset of the recorded neurons. **B.-E.** Quantification of spiking behavior at 7, 14, 17 and 23 DIV. A burst is a sequence of 4 or more spikes with inter-spike-interval lower than 250 μs. Data are shown as mean ± SD; unpaired Student’s t-test (7 DIV, n = 8 neurons; 14 DIV, n = 26 neurons; 17 DIV, n = 32 neurons; 23 DIV, n = 34 neurons). IBI, inter-burst interval. **F.** The AP duration decreases with neuronal maturation. The AP duration was quantified as the full width at half maximal amplitude (FWHM) at 7 DIV (n = 5 neurons, 316 APs), 14 DIV (n = 4, 635 APs) and 17 DIV (n = 5, 1279 APs). Data are shown as mean ± SD; unpaired Student’s t-test. These observations are in agreement with previous work in hippocampal primary cultures (Penn et al 2016). **G.** Under our imaging conditions, we observed virtually no photobleaching of BeRST-1 for the 3 min of continuous illumination with the 635-nm laser at 25 mW/cm^2^ laser power. Top: a representative raw voltage trace. Bottom: the output of the data processing algorithm.

**Figure 1 - figure supplement 2.**
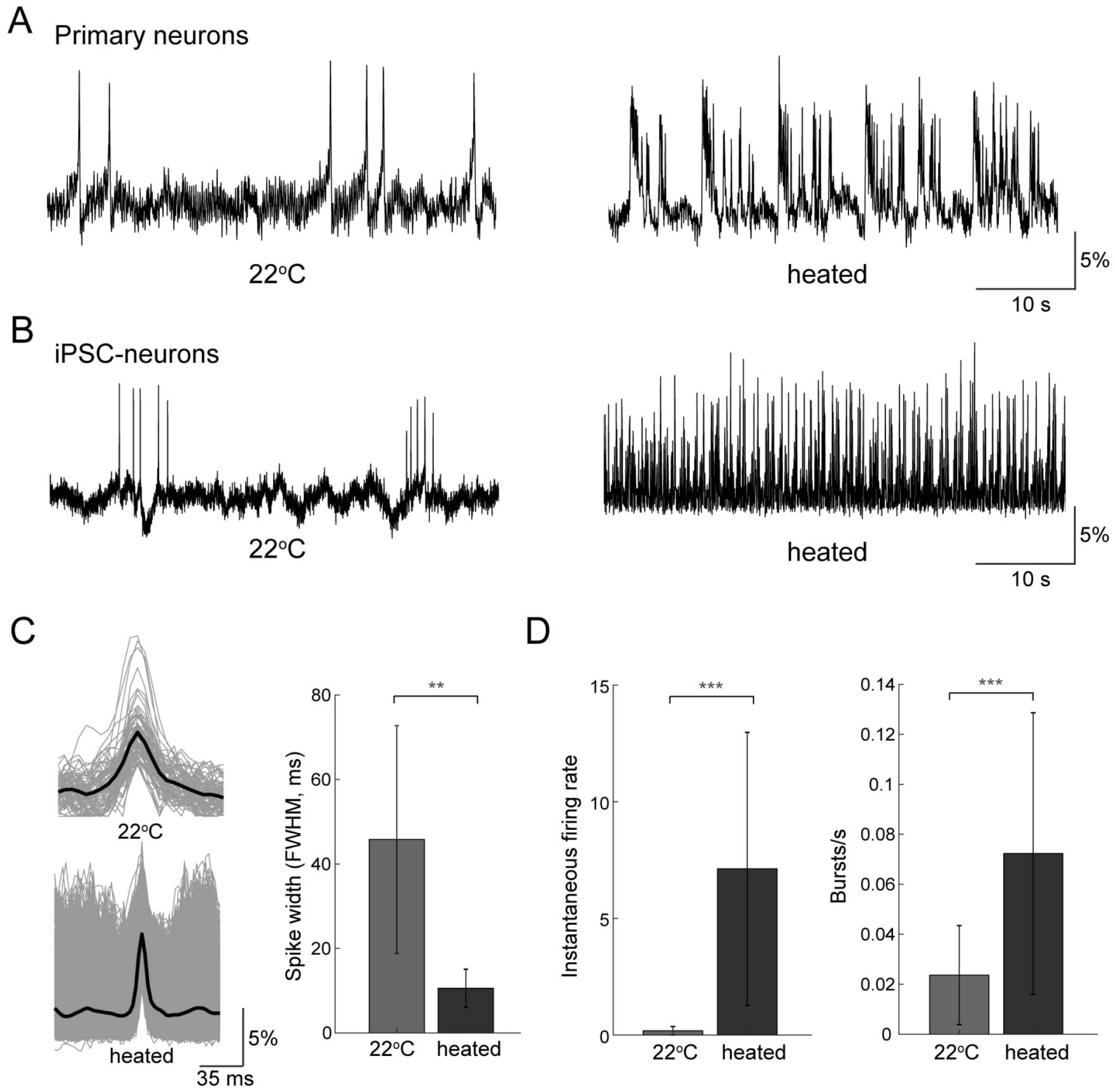
Temperature effects. **A.** Temperature alters intrinsic membrane excitability, response properties and firing rates (Thompson et al 1985). Therefore, careful maintenance of temperature within the imaging chamber was critical for robustness and reproducibility of results. Representative voltage time-courses obtained from a primary neuron at room temperature (22 °C, left) or under perfusion with heated imaging buffer (30 C, right) show a dramatic increase in the firing rate as well as generation of bursts and plateau depolarization. **B.** These effects were also present in hiPCS-derived cultures. Representative voltage time-courses from a human neuron at room temperature (22 °C, left) or under perfusion with heated imaging buffer (32 ° C, right) show a sharp increase in the firing frequency. **C.** In both primary and human neurons, the AP duration was temperature-dependent. For human neurons, the AP duration, quantified as the full width at half maximum amplitude, decreased from 45.8±26.9 to 10.6±4.5 ms with an increase of the bath temperature from 22 °C (n = 11 neurons, 1028 APs) to 32 °C (n=6 neurons, 7000 APs). **D.** Under the same conditions, the instantaneous firing frequency increased from 0.18±0.18 to 7.12±5.85 Hz, and the bursting frequency - from 0.02±0.02 to 0.07±0.05 Hz. Data are shown as mean ± SD; unpaired Student’s t-test.

**Figure 1 - figure supplement 3.**
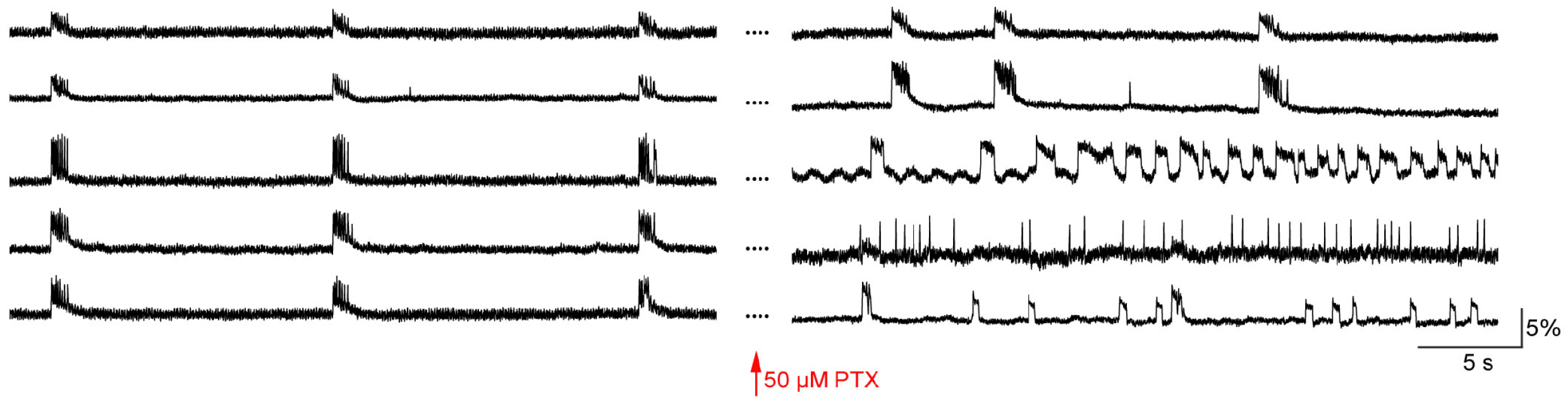
Effect of picrotoxin on spontaneous firing of primary neurons. Perfusion with 50 μM picrotoxin increases the activity in rat hippocampal neurons. Voltage time-courses obtianed from six representative neurons show spontaneous activity prior to pharmacological stimulation (left) and an increase in firing and bursting activity under 50 μM picrotoxin (PTX, right).

**Figure 2 - figure supplement 1.**
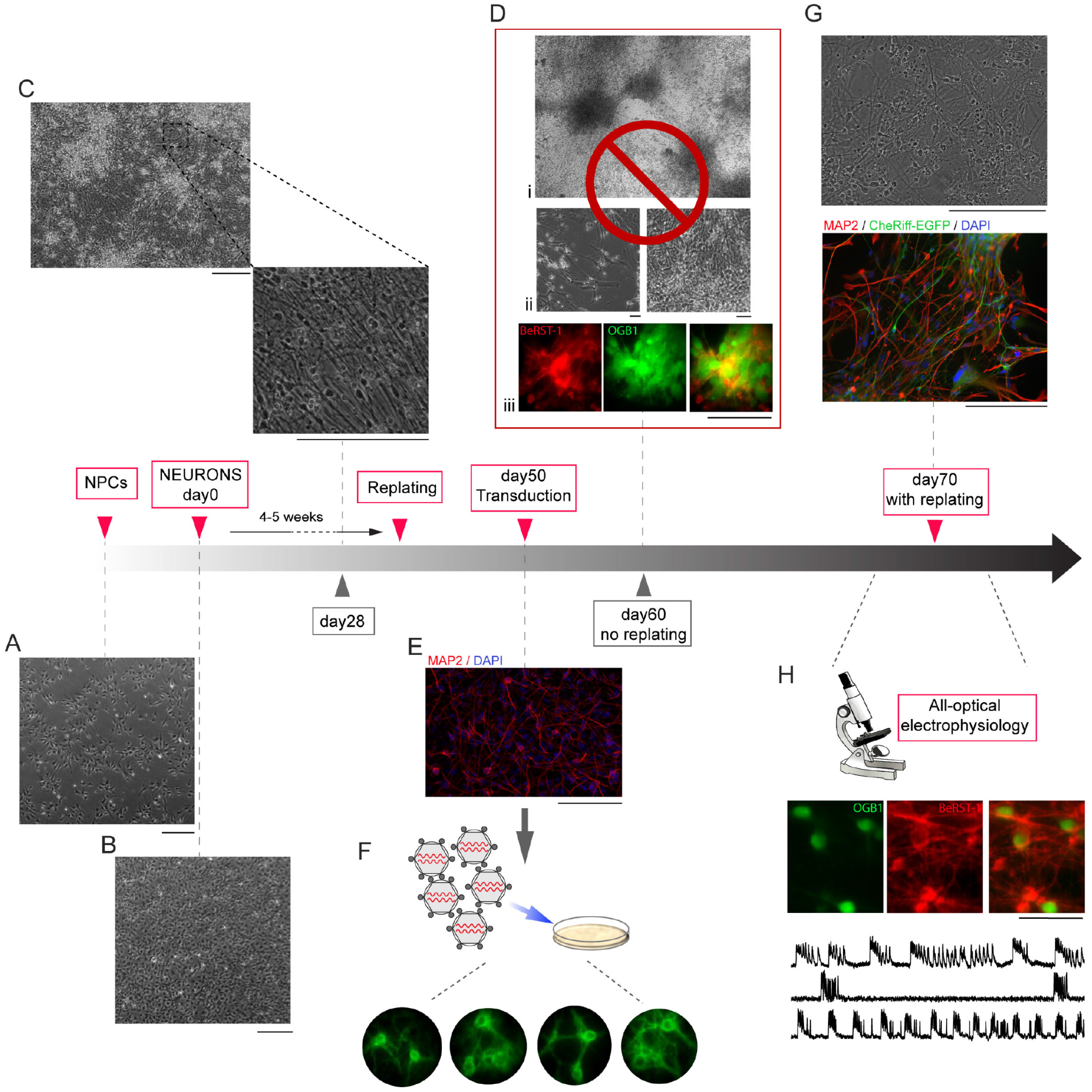
Generation of monolayer cultures of hiPSC-derived neurons

Our goal was to establish a protocol for obtaining healthy/active monolayer and spatially uniform cultures that were critical for the overall success of our experiments. In common practice, cells are typically cultured in high-density to boost the mutual benefit of secreted neurotrophic factors (Yang et al 2017). These high density cultures tend to form large clusters of overlapping neurons. Single-photon fluorescent imaging, however, does not possess depth sectioning. Therefore, this application works best in monolayers of cells, where a possibility for signal crosstalk between adjacent/overlapping neurons is insignificant. Although this cross-talk can be mitigated in part by using oblique illumination (Werley et al 2017) and computational unmixing (Hochbaum et al 2014, Mukamel et al 2009), these remedies are not very effective in overconfluent cultures, in particular when clustered cells fire synchronously. In our hands, high density cultures also exhibited a high rate of cell death due to viral transduction leading to poor expression of the OG actuator CheRiff.

Achieving monolayer cultures with hiPSCs was noticeably more difficult compared to primary cultures due to a well-recognized lengthy process of transition from neural progenitors cells (NPC) to neuronal networks requiring careful control of growth and maturation (Calabrese et al 2019). Specifically, neurons growing for a long period of time in the same dish commonly form clustered, interconnected networks producing robust patterns of network level activity (Segev et al 2001, Wagenaar et al 2006). While this may be desirable for multi-electrode array (MEA) studies (Biffi et al 2013), such networks are suboptimal for single-photon fluorescence imaging due to the abovementioned problem of signal cross-talk. In addition, after more than 5 weeks of growth in the same dish, we observed that hiPSC-derived cell cultures had the following 3 properties aversive for optical imaging:

i. They easily detached from the substrate when handled.
ii. They often contained a layer of dead cells attached to healthy neurons. Dead cells sequestered optical probes generating strong background fluorescence.
iii. They contained NPCs that remained undifferentiated, continued to divide and, in many cases, dominated over differentiated neurons competing with them for the growth substrate. Under these conditions, NPCs could also deprive differentiated neurons of their nutrients.

To mitigate these problems, we modified a published cell culture protocol (Calabrese et al 2019) resulting in well-adhered, relatively sparse and uniform cultures with limited cell clustering. Our protocol included the following critical steps:

i. The initial 4-5 weeks of NPC differentiation in 10-cm Petri dishes.
ii. Replating the cells into 35-mm imaging plates only after mature neurons with robust and long interconnections were formed.
iii. Further 2-3 weeks of differentiation to allow recovery, re-growth of connections and neuronal maturation.
iv. Transduction with lentivirus carrying CheRiff-EGFP expression cassette.
v. All-optical electrophysiology performed 7 to 10 days after transfection when robust

EGFP fluorescence indicated successful expression of the transgene.

Depending on the cell line, the maturation time-course varied. Therefore, the timing of replating and transfection was determined based on observation of the culture growth and connectivity. Best results were achieved when replating occurred prior to slowing down of growth and connectivity and a decrease in cell viability.

**A.** Transmitted light image of NPCs (scale bar, 200 μm) plated on 10-cm Petri dishes.

**B.** Transmitted light image of NPCs culture that reached a confluency of 90%, which is the point when bFGF was withdrawn to start differentiation. Scale bar, 200 μm.

**C.** Transmitted light image of developing human neurons after 4 weeks of differentiation with long and robust connections. Scale bar, 200 μm.

**D.** hiPSC-derived neuronal cultures after ~8 weeks of differentiation in 10-cm Petri dishes, without replating. Transmitted light images show formation of clusters (i), detached cells (ii, left) and clumps of cells on top of each other (ii, right). Clumps of cells from human neurons, which have been differentiating for 8 weeks, stained with BeRST-1 and loaded with OGB1 (iii). Scale bars, 50 μm.

**E.** Replating into 35-mm imaging plates at week 4-5 of differentiation allows formation of homogenous and sparse neuronal cultures. The culture was immune-stained with MAP2 (red), the nuclei were counterstained with DAPI (blue). Scale bar, 50 μm.

**F.** Monolayer cultures facilitate expression of CheRiff-EGFP via lentiviral transduction. The epifluorescence images show representative examples of CheRiff-EGFP-expressing neurons 10 days after the viral transfection.

**G.** Human neurons after 10 weeks of differentiation, which were replated at week 4. The transmitted light image shows a homogenous, monolayer culture cells with no clustering. The immunostaining image indicates mature neurons expressing MAP2 (red) and CheRiff-EGFP (green). The nuclei were counterstained with DAPI (blue). Scale bar, 200 μm.

**H.** hiPSC neurons were imaged between week 9 and 10 of differentiation. Top: Epifluorescence images of cells loaded with OGB1 and stained with BeRST-1. Bottom: three representative BeRST-1 traces of spontaneous activity.

## Acknowledgements

We thank Eran Mukamel for helpful discussions on signal analysis and gratefully acknowledge support from the NIH BRAIN Initiative (R21EY030727, R01MH111359, U01NS094232); NIH R01NS098088; NIH U19MH1073671, part of the National Cooperative Reprogrammed Cell Research Groups (NCRCRG) to Study Mental Illness; Research Council of Norway (223273, 248828, 283798); K.G. Jebsen Stiftelsen and South-Eastern Norway Regional Health Authority (#2018094). C.A.T. was partly funded by K01 NIAAA (K01AA026911) and a CDKL5 Loulou Foundation grant.

## References

Bando Y, Sakamoto M, Kim S, Ayzenshtat I, Yuste R. 2019. Comparative Evaluation of Genetically Encoded Voltage Indicators. Cell Rep 26: 802–13 e4

Bean BP. 2007. The action potential in mammalian central neurons. Nat Rev Neurosci 8: 451–65

Biffi E, Regalia G, Menegon A, Ferrigno G, Pedrocchi A. 2013. The influence of neuronal density and maturation on network activity of hippocampal cell cultures: a methodological study. PLoS One 8: e83899

Brennand KJ, Simone A, Jou J, Gelboin-Burkhart C, Tran N, et al. 2011. Modelling schizophrenia using human induced pluripotent stem cells. Nature 473: 221–5

Calabrese B, Powers RM, Slepian AJ, Halpain S. 2019. Post-differentiation Replating of Human Pluripotent Stem Cell-derived Neurons for High-content Screening of Neuritogenesis and Synapse Maturation. J Vis Exp

Chen L, Deng Y, Luo W, Wang Z, Zeng S. Detection of bursts in neuronal spike trains by the mean interspike interval method. Progress in Natural Science 19: 229–35

Deisseroth K. 2015. Optogenetics: 10 years of microbial opsins in neuroscience. Nat Neurosci 18: 1213–25

Detrait ER, Bowers WJ, Halterman MW, Giuliano RE, Bennice L, et al. 2002. Reporter gene transfer induces apoptosis in primary cortical neurons. Mol Ther 5: 723–30

Ding B, Kilpatrick DL. 2013. Lentiviral vector production, titration, and transduction of primary neurons. Methods Mol Biol 1018: 119–31

Emiliani V, Cohen AE, Deisseroth K, Hausser M. 2015. All-Optical Interrogation of Neural Circuits. J Neurosci 35: 13917–26

Goslin K, Asmussen H, Banker G. 1991. Rat hippocampal neurons in low-density culture In Culturing Nerve Cells, pp. 339–70: MIT Press

Griesi-Oliveira K, Acab A, Gupta AR, Sunaga DY, Chailangkarn T, et al. 2015. Modeling non-syndromic autism and the impact of TRPC6 disruption in human neurons. Mol Psychiatry 20: 1350–65

Hanson HH, Reilly JE, Lee R, Janssen WG, Phillips GR. 2010. Streamlined embedding of cell monolayers on gridded glass-bottom imaging dishes for correlative light and electron microscopy. Microsc Microanal 16: 747–54

Hochbaum DR, Zhao Y, Farhi SL, Klapoetke N, Werley CA, et al. 2014. All-optical electrophysiology in mammalian neurons using engineered microbial rhodopsins. Nat Methods 11: 825–33

Huang YL, Walker AS, Miller EW. 2015. A Photostable Silicon Rhodamine Platform for Optical Voltage Sensing. J Am Chem Soc 137: 10767–76

Iakoucheva LM, Muotri AR, Sebat J. 2019. Getting to the Cores of Autism. Cell 178: 1287–98

Johnston S, Parylak S, Kim S, Mac N, Lim C, et al. 2020. AAV Ablates Neurogenesis in the Adult Murine Hippocampus. bioRxiv

Kiskinis E, Kralj JM, Zou P, Weinstein EN, Zhang H, et al. 2018. All-Optical Electrophysiology for High-Throughput Functional Characterization of a Human iPSC-Derived Motor Neuron Model of ALS. Stem Cell Reports 10: 1991–2004

Kulkarni RU, Miller EW. 2017. Voltage Imaging: Pitfalls and Potential. Biochemistry 56: 5171–77

Marchetto MC, Belinson H, Tian Y, Freitas BC, Fu C, et al. 2017. Altered proliferation and networks in neural cells derived from idiopathic autistic individuals. Mol Psychiatry 22: 820–35

Marchetto MC, Carromeu C, Acab A, Yu D, Yeo GW, et al. 2010. A model for neural development and treatment of Rett syndrome using human induced pluripotent stem cells. Cell 143: 527–39

Mertens J, Wang QW, Kim Y, Yu DX, Pham S, et al. 2015. Differential responses to lithium in hyperexcitable neurons from patients with bipolar disorder. Nature 527: 95–9

Miller EW. 2016. Small molecule fluorescent voltage indicators for studying membrane potential. Curr Opin Chem Biol 33: 74–80

Mukamel EA, Nimmerjahn A, Schnitzer MJ. 2009. Automated analysis of cellular signals from large-scale calcium imaging data. Neuron 63: 747–60

Mukhopadhyay S, Ray GC. 1998. A new interpretation of nonlinear energy operator and its efficacy in spike detection. IEEE Trans Biomed Eng 45: 180–7

Paavilainen T, Pelkonen A, Makinen ME, Peltola M, Huhtala H, et al. 2018. Effect of prolonged differentiation on functional maturation of human pluripotent stem cell-derived neuronal cultures. Stem Cell Res 27: 151–61

Penn Y, Segal M, Moses E. 2016. Network synchronization in hippocampal neurons. Proc Natl Acad Sci U SA 113:3341–6

Peterka DS, Takahashi H, Yuste R. 2011. Imaging voltage in neurons. Neuron 69: 9–21

Segev R, Shapira Y, Benveniste M, Ben-Jacob E. 2001. Observations and modeling of synchronized bursting in two-dimensional neural networks. Phys Rev E Stat Nonlin Soft Matter Phys 64: 011920

Shi Y, Kirwan P, Smith J, Robinson HP, Livesey FJ. 2012. Human cerebral cortex development from pluripotent stem cells to functional excitatory synapses. Nat Neurosci 15: 477–86, S1

Takahashi K, Tanabe K, Ohnuki M, Narita M, Ichisaka T, et al. 2007. Induction of pluripotent stem cells from adult human fibroblasts by defined factors. Cell 131: 861–72

Thompson SM, Masukawa LM, Prince DA. 1985. Temperature dependence of intrinsic membrane properties and synaptic potentials in hippocampal CA1 neurons in vitro. J Neurosci 5: 817–24

Trainito C, von Nicolai C, Miller EK, Siegel M. 2019. Extracellular Spike Waveform Dissociates Four Functionally Distinct Cell Classes in Primate Cortex. Curr Biol 29: 2973–82 e5

Wagenaar DA, Pine J, Potter SM. 2006. An extremely rich repertoire of bursting patterns during the development of cortical cultures. BMC Neurosci 7: 11

Walker AS, Raliski BK, Karbasi K, Zhang P, Sanders K, Miller EW. 2020a. Optical spike detection and connectivity analysis with a far-red voltage-sensitive fluorophore reveals changes to network connectivity in development and disease. bioRxiv

Walker AS, Raliski BK, Nguyen DV, Zhang P, Sanders K, et al. 2020b. Imaging voltage in complete neuronal networks within patterned microislands reveals preferential wiring of excitatory hippocampal neurons. bioRxiv

Werley CA, Brookings T, Upadhyay H, Williams LA, McManus OB, Dempsey GT. 2017. All-Optical Electrophysiology for Disease Modeling and Pharmacological Characterization of Neurons. Curr Protoc Pharmacol 78: 11 20 1–11 20 24

Yang HH, St-Pierre F. 2016. Genetically Encoded Voltage Indicators: Opportunities and Challenges. J Neurosci 36: 9977–89

Yang Q, Ke Y, Luo J, Tang Y. 2017. Protocol for culturing low density pure rat hippocampal neurons supported by mature mixed neuron cultures. J Neurosci Methods 277: 38–45

Yu J, Vodyanik MA, Smuga-Otto K, Antosiewicz-Bourget J, Frane JL, et al. 2007. Induced pluripotent stem cell lines derived from human somatic cells. Science 318: 1917–20

Zhang ZN, Freitas BC, Qian H, Lux J, Acab A, et al. 2016. Layered hydrogels accelerate iPSC-derived neuronal maturation and reveal migration defects caused by MeCP2 dysfunction. Proc Natl Acad Sci US A 113: 3185–90

